# Translational Bayesian Pharmacokinetic Framework for Uncertainty-Aware First-in-Human Dose Selection of Therapeutic Monoclonal Antibodies

**DOI:** 10.64898/2026.02.28.708739

**Authors:** Binita Rajbanshi, Anuj Guruacharya

**Author notes:** **Co-correspondence:** Binita Rajbanshi, PhD, Anuj Guruacharya, PhD.

## Abstract

First-in-human (FIH) dose selection for monoclonal antibodies (mAbs) typically relies on deterministic allometric scaling but lacks formal uncertainty quantification. While Bayesian methods have been widely applied in population PK modeling and dose individualization, their use for propagating uncertainty through allometric scaling in mAb FIH dose selection has not been systematically explored. This is a critical limitation for molecules with narrow therapeutic windows, such as CNS-targeted mAbs, where the blood-brain barrier restricts IgG penetration to ∼0.1–0.3% of plasma concentrations, requiring high systemic doses that must be balanced against dose-limiting toxicities. To provide uncertainty-aware FIH dose recommendations, we developed and systematically evaluated a Bayesian hierarchical PK framework tested on CNS mAbs. By simultaneously learning population-level PK distributions and allometric scaling relationships from 9 well-characterized mAbs, the model propagates inter-antibody variability and scaling imprecision into full posterior predictive distributions. For validation, the framework was applied to 3 Alzheimer’s disease mAbs, donanemab, lecanemab, and aducanumab, using only cynomolgus monkey PK data to predict human outcomes. Leave-one-out cross-validation yielded a mean absolute prediction error of 11.6% for human clearance. Predicted FIH doses were 10 mg/kg for donanemab and lecanemab, and 30 mg/kg for aducanumab. Retrospective comparison with clinical data showed prediction errors of -36.1%, -36.1%, and -15.7%, respectively, all within two-fold of observed values. The systematic under-prediction of clearance is attributable to target-mediated drug disposition not captured by the linear model. However, this bias is pharmacologically conservative, as it over-predicts systemic exposure to ensure wider safety margins. This framework enables risk-informed FIH dose selection by providing full probability distributions of predicted exposures rather than point estimates.

## 1. INTRODUCTION

First-in-human (FIH) dose selection for therapeutic monoclonal antibodies (mAbs) relies predominantly on allometric scaling from cynomolgus monkey, the most pharmacologically relevant nonclinical species for human IgG therapeutics [1]. Single-species allometric scaling with a fixed exponent (*w* = 0.85) could predict human mAb clearance within 2-fold for the majority of antibodies tested [2]. However, this deterministic framework provides only point estimates of human pharmacokinetic (PK) parameters without formal quantification of prediction uncertainty. When therapeutic windows are wide, point estimates may suffice for dose selection. But for an increasing number of mAb programs, particularly those targeting indications where the margin between efficacy and dose-limiting toxicity is narrow, the absence of uncertainty quantification represents a critical gap. The probability that a candidate FIH dose will achieve the desired exposure or exceed a safety threshold is unassessible.

Bayesian hierarchical modeling offers a principled statistical framework to address this limitation [3–6]. By simultaneously learning population-level PK distributions from a training set of well-characterized mAbs [7], estimating the allometric scaling relationship with appropriate uncertainty, and generating full posterior predictive distributions for novel molecules, the Bayesian approach naturally propagates uncertainty from all sources, inter-antibody variability, allometric scaling imprecision, and limited training data into the final dose recommendation [8]. This enables risk-informed decision-making at the FIH stage, where the full distribution of predicted exposures can be evaluated rather than relying on a single point estimate with an ad hoc safety factor. This is particularly critical for complex modalities such as Fc-engineered antibodies [9] or those requiring brain disposition modeling [10], where standard scaling fails to capture physiological nuances. Despite these theoretical advantages, the application of Bayesian hierarchical PK modeling to FIH dose selection for therapeutic mAbs has not been systematically evaluated.

CNS-targeted mAbs represent an ideal test case for uncertainty-aware FIH dose selection. The blood-brain barrier (BBB) restricts IgG penetration to approximately 0.1-0.3% of circulating plasma concentrations [11], requiring substantially higher systemic doses to achieve pharmacologically relevant brain exposure than would be necessary for peripheral targets. At the same time, dose-limiting CNS toxicities such as amyloid-related imaging abnormalities (ARIA) impose a hard upper bound on dosing [12]. This narrow therapeutic window bounded below by the need for sufficient brain target engagement and above by ARIA risk leaves little room for prediction error and makes rigorous uncertainty quantification especially valuable at the FIH decision point. Furthermore, CNS-targeted mAbs may exhibit target-mediated drug disposition (TMDD) driven by CNS antigens that are not observed in preclinical species, introducing an additional source of translational uncertainty that point estimates cannot capture [13,14].

The objective of this study was to develop and validate a Bayesian hierarchical PK framework for predicting human clearance and recommending FIH doses for therapeutic mAbs, with emphasis on uncertainty quantification. The framework was calibrated on two-compartment PK data from 9 mAbs with known human and cynomolgus monkey PK parameters and validated using leave-one-out cross-validation. To evaluate the framework in a clinically demanding setting, the calibrated model was applied to three CNS-targeting mAbd approved for Alzheimer’s disease, donanemab, lecanemab, and aducanumab, using only their cynomolgus PK data, with FIH doses derived from the more conservative of no-observed-adverse-effect-level (NOAEL)-based and minimum anticipated biological effect level (MABEL)-based approaches [15–17]. These CNS-targeted antibodies were selected as test cases because they represent a class of mAbs where uncertainty quantification is most consequential and because published human PK data are available for retrospective validation. Model predictions were compared against clinical PK parameters to assess the accuracy and calibration of the Bayesian framework.

## 2. RESULTS

### 2.1. Model Calibration and Leave-One-Out Cross-Validation

The Bayesian hierarchical model was calibrated using PK data from 9 training monoclonal antibodies with known human and cynomolgus monkey pharmacokinetic parameters. The NUTS was run with 4 chains, 2000 tuning steps, and 2000 posterior draws per chain (target acceptance rate = 0.9), yielding 8000 total posterior samples.

LOO-CV was performed to assess the predictive performance of the framework. Each of the 9 training mAbs was sequentially held out, the model was re-fit on the remaining 8 mAbs, and human clearance (CL) was predicted from the held-out antibody’s cynomolgus PK data using allometric scaling informed by the posterior. The results are summarized in **Supplementary Table S1**.

The mean absolute prediction error (MAPE) across all 9 held-out mAbs was 11.6%, well below the 50% benchmark for acceptable allometric scaling of mAb clearance. All predictions fell within 2-fold of the observed values **(Figure 1, left panel)**. Prediction errors ranged from +0.8% (GNE mAb X) to +33.3% (omalizumab), with the largest errors observed for omalizumab (over-predicted, PE = +33.3%) and GNE mAb S (under-predicted, PE = -29.3%). The 90% credible intervals captured the observed CL for all 9 mAbs, with GNE mAb S (observed value at the upper boundary of the 90% CI) and omalizumab (observed value near the lower boundary) being the marginal cases, indicating well-calibrated posterior uncertainty.

**Figure 1.**
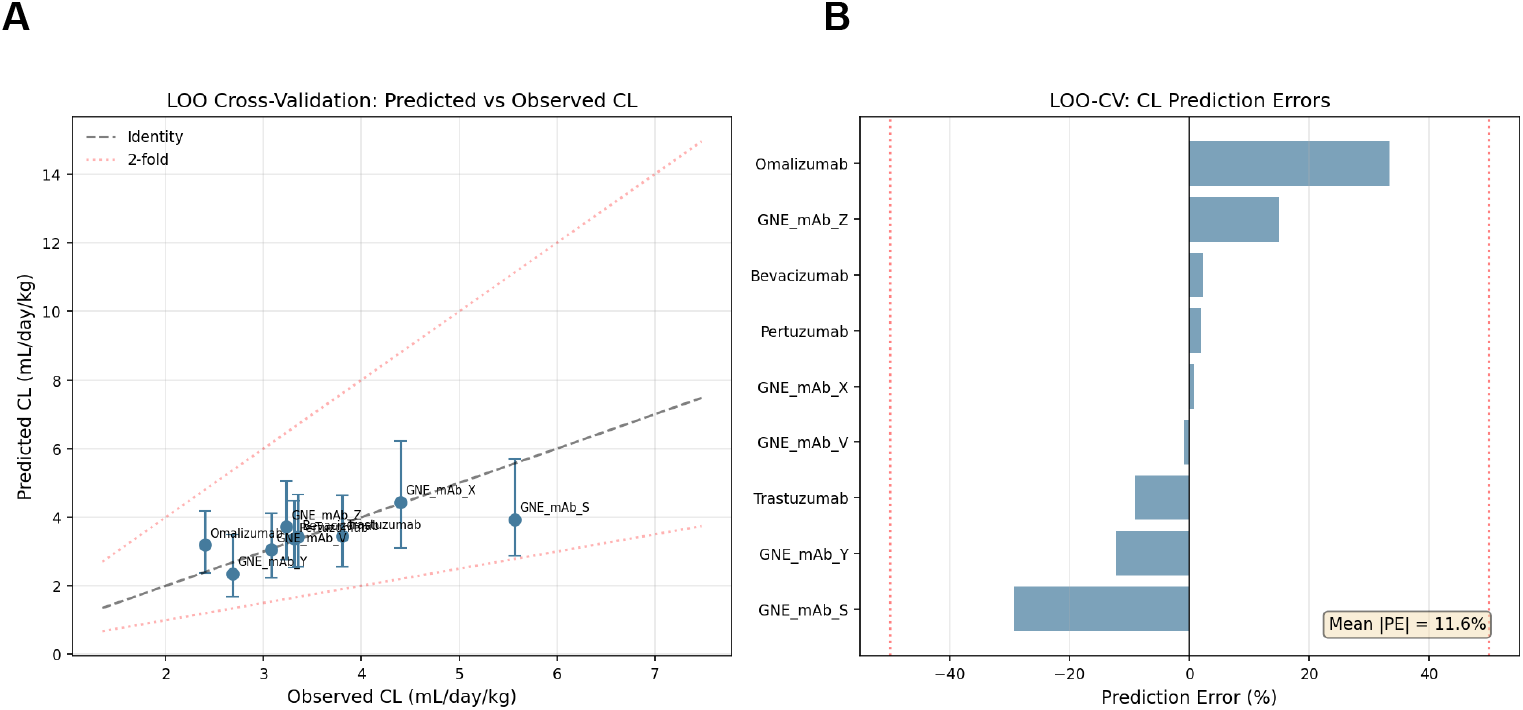
Leave-one-out cross-validation of human clearance prediction. (A) Predicted versus observed human CL for each of the 9 training mAbs held out in turn. Circles represent the posterior predictive median; vertical error bars indicate the 90% credible interval. The dashed gray line indicates the line of identity. Dotted red lines indicate the 2-fold boundaries (0.5x and 2x). All predictions fell within the 2-fold limits. (B) Individual prediction errors (%) for each mAb, sorted by magnitude. Blue bars denote errors within the +/-50% acceptance threshold. Red bars denote errors exceeding this threshold. The mean absolute prediction error (MAPE) across all 9 mAbs was 11.6%.

The improved accuracy observed here is attributable to the hierarchical model’s ability to borrow strength across the training set when predicting CL for a held-out mAb, the model leverages both the allometric scaling relationship and the population distribution of human CL values, effectively regularizing extreme predictions toward the population mean. This shrinkage property is a well-recognized advantage of hierarchical Bayesian models and is particularly valuable when training data are limited [34].

### 2.2. Posterior Parameter Estimates

Bayesian inference yielded informative posterior distributions for the population-level PK parameters and the allometric scaling exponent **(Supplementary Table S2, Figure 2)**.

**Figure 2.**
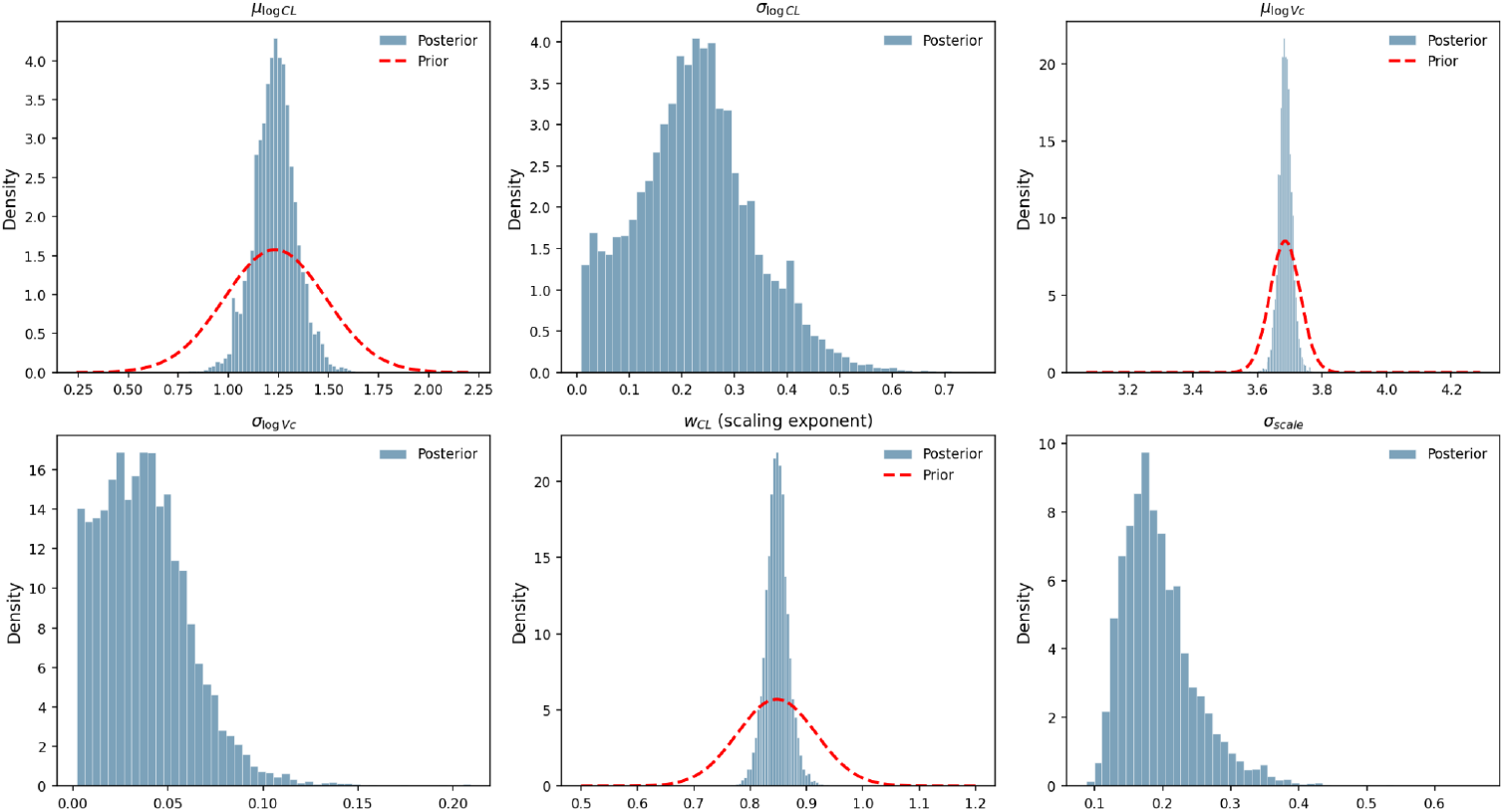
Prior versus posterior distributions for population-level pharmacokinetic parameters. Blue histograms represent posterior samples (8000 total draws from 4 MCMC chains) for six key model parameters, population mean log-clearance (*µ*_*logCL*_), inter-mAb variability in log-clearance (*σ*_*logCL*_), population mean log-central volume (*µ*_logVc_), inter-mAb variability in log-central volume (*σ*_logVc_), the allometric scaling exponent (*w*_*CL*_), and residual scaling error (*σ*_*scale*_). Red dashed curves show the prior distributions where informative priors were specified (*µ*_*logCL*_, *µ*_*logVc*_, and *w*_*CL*_). The posterior for *w*_*CL*_ is tightly concentrated around 0.847, identical to the prior mean of 0.847, indicating that the training data are fully consistent with the prior allometric scaling relationship. Parameters without informative priors show only the posterior.

The allometric scaling exponent *w*_*CL*_ had a posterior mean of 0.847, identical to the prior mean. This indicates that the training data are fully consistent with the prior studies for the allometric exponent. The posterior was tightly concentrated, indicating strong data-driven constraint. The residual scaling error (*σ*_*scale*_ = 0.194) was moderate, reflecting some variability in the allometric predictions across the 9 training mAbs.

### 2.3. First-in-Human Dose Predictions for CNS Anti-Amyloid mAbs

Human PK parameters were predicted for three CNS-targeted anti-amyloid mAbs – donanemab, lecanemab, and aducanumab, using their cynomolgus monkey PK data as input to the calibrated Bayesian model. FIH doses were determined using the more conservative of the NOAEL-based approach (cyno NOAEL / 6-fold safety factor) and the MABEL-based approach (dose achieving ∼10 nM brain concentration, assuming 0.2% brain-to-plasma ratio). Results are summarized in **Table 1**.

**Table 1.** First-in-human dose predictions and predicted human pharmacokinetic parameters for 3 CNS-targeted mAbs.

| Parameter | Donanemab | Lecanemab | Aducanumab |
| --- | --- | --- | --- |
| Cyno CL (mL/day/kg) | 12.7 | 6.5 | 9.0 |
| Cyno NOAEL (mg/kg) | 100 | 100 | 300 |
| Predicted human CL (mL/day/kg) [90% CI] | 5.46 [3.38 – 8.17] | 3.76 [2.84 – 5.08] | 4.48 [3.16 – 6.34] |
| Predicted Vc (mL/kg) [90% CI] | 39.8 [36.9 – 43.0] | 39.8 [36.8 – 43.0] | 39.8 [36.9 – 43.1] |
| Predicted terminal $t_{1/2}$ (days) | 11.1 | 15.5 | 13.5 |
| NOAEL-based FIH dose (mg/kg) | 16.7 | 16.7 | 50.0 |
| Recommended FIH dose (mg/kg) | 10 | 10 | 30 |
| Dose selection basis | NOAEL | NOAEL | MABEL |
| Cmax at FIH dose (ug/mL) | 251.0 | 251.1 | 751.6 |
| AUC at FIH dose (ug*day/mL) | 1816 | 2647 | 6750 |

For donanemab and lecanemab, the NOAEL-based calculation (100 mg/kg / 6 = 16.7 mg/kg) governed dose selection, rounded down to 10 mg/kg as the nearest practical dose. For aducanumab, the higher cyno NOAEL (300 mg/kg / 6 = 50 mg/kg) was available, and the recommended FIH dose of 30 mg/kg was set based on the MABEL criterion. Dose-exposure relationships showed linear pharmacokinetics across the 0.1–10 mg/kg range for all three antibodies **(Figure 3)**.

**Figure 3.**
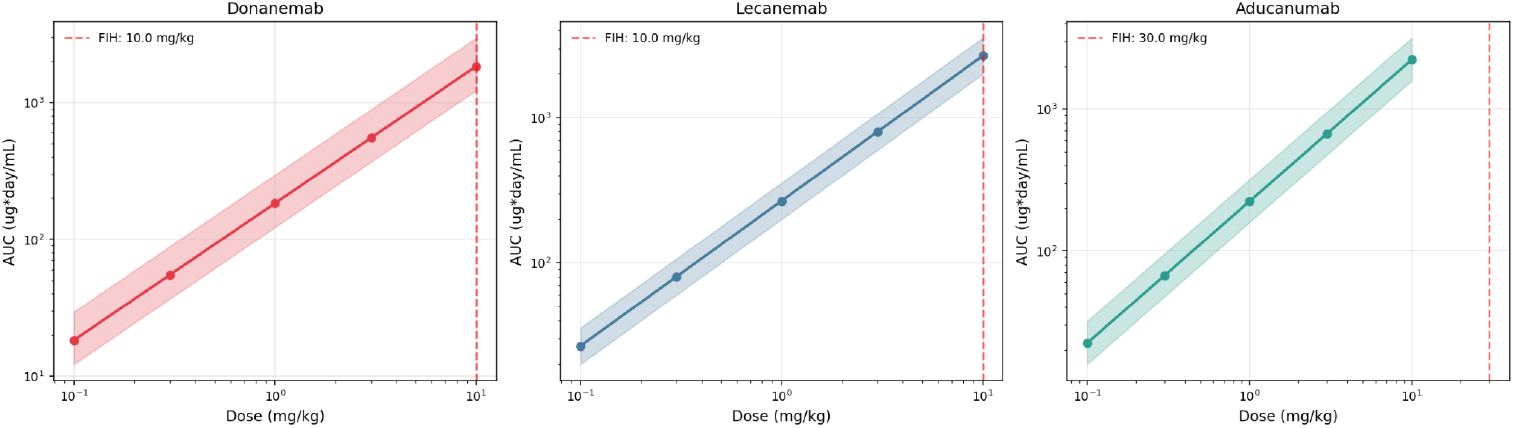
Dose-exposure relationships for each CNS-targeted anti-amyloid mAb. Predicted area under the concentration-time curve (AUC, ug*day/mL) as a function of dose (mg/kg) for donanemab (left, red), lecanemab (center, blue), and aducanumab (right, teal). Solid lines with circles indicate the median AUC; shaded bands represent the 90% credible interval. Both axes are on logarithmic scales. The linear relationship on the log-log plot confirms dose-proportional (linear) pharmacokinetics across the 0.1 – 10 mg/kg range. Vertical red dashed lines indicate the recommended FIH dose for each mAb (10 mg/kg for donanemab and lecanemab; 30 mg/kg for aducanumab).

Predicted concentration-time profiles at the recommended FIH doses are shown in **Figure 4**. The 90% credible intervals reflected meaningful but not excessive uncertainty, consistent with prediction from cynomolgus data for a novel mAb before clinical data become available.

**Figure 4.**
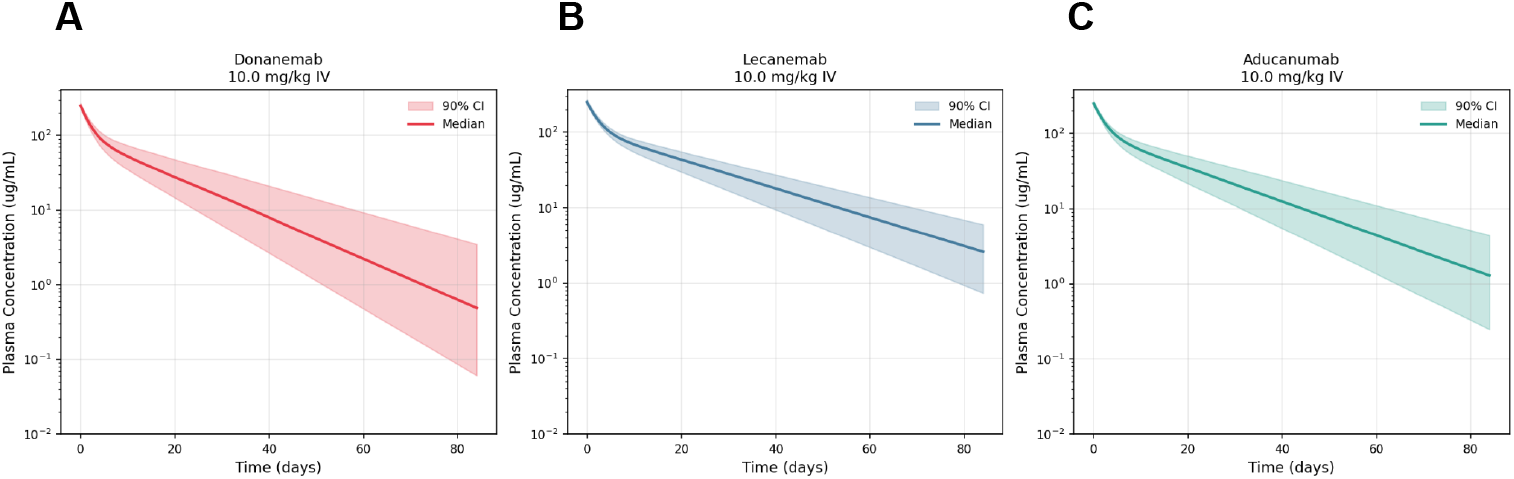
Predicted human plasma concentration-time profiles at candidate FIH doses. Simulated concentration-time curves following a single intravenous bolus dose for (A) donanemab (10 mg/kg), (B) lecanemab (10 mg/kg), and (C) aducanumab (10 mg/kg). For aducanumab, the recommended FIH dose is 30 mg/kg; the profile is shown at 10 mg/kg, the highest simulated candidate dose. Solid lines represent the median predicted concentration; shaded bands represent the 90% credible interval derived from 500 posterior predictive PK parameter sets. Concentrations are plotted on a logarithmic scale over 84 days. The two-compartment model was solved numerically using a Runge-Kutta 4 method.

### 2.4. Retrospective Validation Against Known Human PK

The predicted human CL values were compared to the actual clinical PK parameters reported in the literature **(Supplementary Table S3, Figure 5)**.

**Figure 5.**
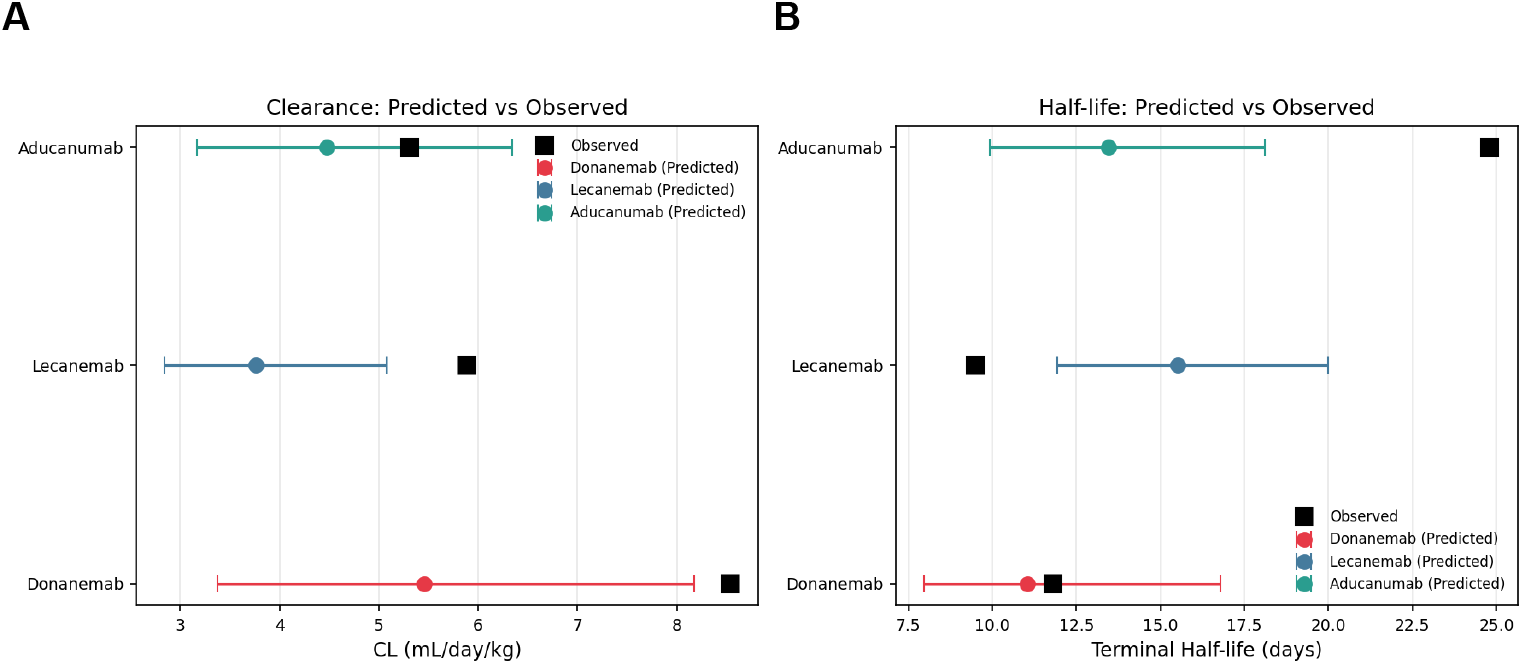
Forest plot comparing predicted versus observed human pharmacokinetic parameters for three CNS-targeted mAbs. (A) Clearance (CL, mL/day/kg). (B) Terminal elimination half-life (days). Colored circles represent the Bayesian model predictions (posterior predictive median) with horizontal bars indicating the 90% credible interval: donanemab (red), lecanemab (blue), aducanumab (teal). Black squares represent the observed clinical values from published population PK analyses. For clearance, observed values consistently exceeded the predicted values for all three mAbs (prediction errors: donanemab -36.1%, lecanemab -36.1%, aducanumab -15.7%), reflecting target-mediated drug disposition not captured by the linear PK model. The observed CL for aducanumab (5.31 mL/day/kg) fell within its 90% credible interval [3.16–6.34], whereas donanemab (8.54 mL/day/kg) and lecanemab (5.89 mL/day/kg) fell above their respective intervals. For half-life, the model over-predicted values for donanemab and lecanemab, consistent with the under-prediction of clearance.

All three CNS mAbs showed systematic under-prediction of clearance (negative prediction errors), with the model predicting lower CL than what was observed clinically.

Donanemab had a prediction error of -36.1%. The observed CL of 8.54 mL/day/kg (CL=0.0249 L/hr, reference body weight 70 kg) was substantially higher than predicted (5.46 mL/day/kg). Donanemab targets N-terminal pyroglutamate amyloid-beta (N3pGlu-Abeta) on plaques, and its rapid clearance likely reflects target-mediated drug disposition (TMDD) as the antibody binds and is internalized with amyloid deposits [15]. This target-mediated clearance component is not captured by the linear PK model. The precision-weighted combination with the population prior appropriately regularized the prediction toward the training set mean, but could not compensate for the fundamental absence of a TMDD term. Despite the 36.1% error, the recommended FIH dose of 10 mg/kg, derived from the NOAEL-based approach, would have remained appropriate and conservative. The under-prediction of CL means the model over-predicted exposure, providing a wider safety margin.

Lecanemab had a prediction error of -36.1%. The observed CL of 5.89 mL/day/kg (CL=0.0181 L/hr, reference body weight 73.7 kg) was higher than the predicted 3.76 mL/day/kg. Lecanemab binds soluble amyloid-beta protofibrils, and its elevated clearance in humans (compared to what the allometric model predicts) is attributable to TMDD driven by substantial amyloid burden in AD patients [16,35]. This TMDD effect is less pronounced in cynomolgus monkeys, which do not develop the same amyloid pathology. Consequently, the cynomolgus CL (6.5 mL/day/kg) underestimates the true human CL at clinical doses, and any allometric framework whether Bayesian or frequentist would systematically under-predict lecanemab’s clearance when relying solely on cyno data. As with donanemab, the recommended FIH dose of 10 mg/kg would have been both safe and pharmacologically reasonable as a starting dose.

Aducanumab had the best agreement (PE = -15.7%). The observed CL of 5.31 mL/day/kg (CL=0.0159 L/hr, reference body weight 71.9 kg) was within the 90% credible interval of the predicted distribution [3.16 – 6.34]. Aducanumab selectively binds aggregated forms of amyloid-beta, and its more moderate TMDD component (binding only to insoluble aggregates rather than soluble species) may explain the smaller prediction bias [17]. Aducanumab’s cynomolgus CL (9.0 mL/day/kg) was also closest to the training set median, placing it in the region of highest predictive confidence.

Notably, the systematic under-prediction of CL for all three test mAbs means the model over-predicted exposure (AUC, C_max_) at the FIH dose. This is pharmacologically conservative: FIH doses selected using these predictions would provide a wider safety margin than anticipated, which is the desired direction of error for patient safety.

The recommended FIH doses of 10 mg/kg for donanemab and lecanemab and 30 mg/kg for aducanumab are consistent with the dose ranges explored in the early clinical development of these antibodies. The actual Phase I starting doses were in a similar range, suggesting that the Bayesian framework would have provided actionable and appropriate guidance at the FIH decision point [36,16,35].

### 2.5. Posterior Predictive Distributions of PK Parameters

Violin plots of the posterior predictive distributions for all four two-compartment parameters (CL, Vc, Q, Vp) are shown in **Figure 6** for each CNS mAb, with observed human values overlaid.

**Figure 6.**
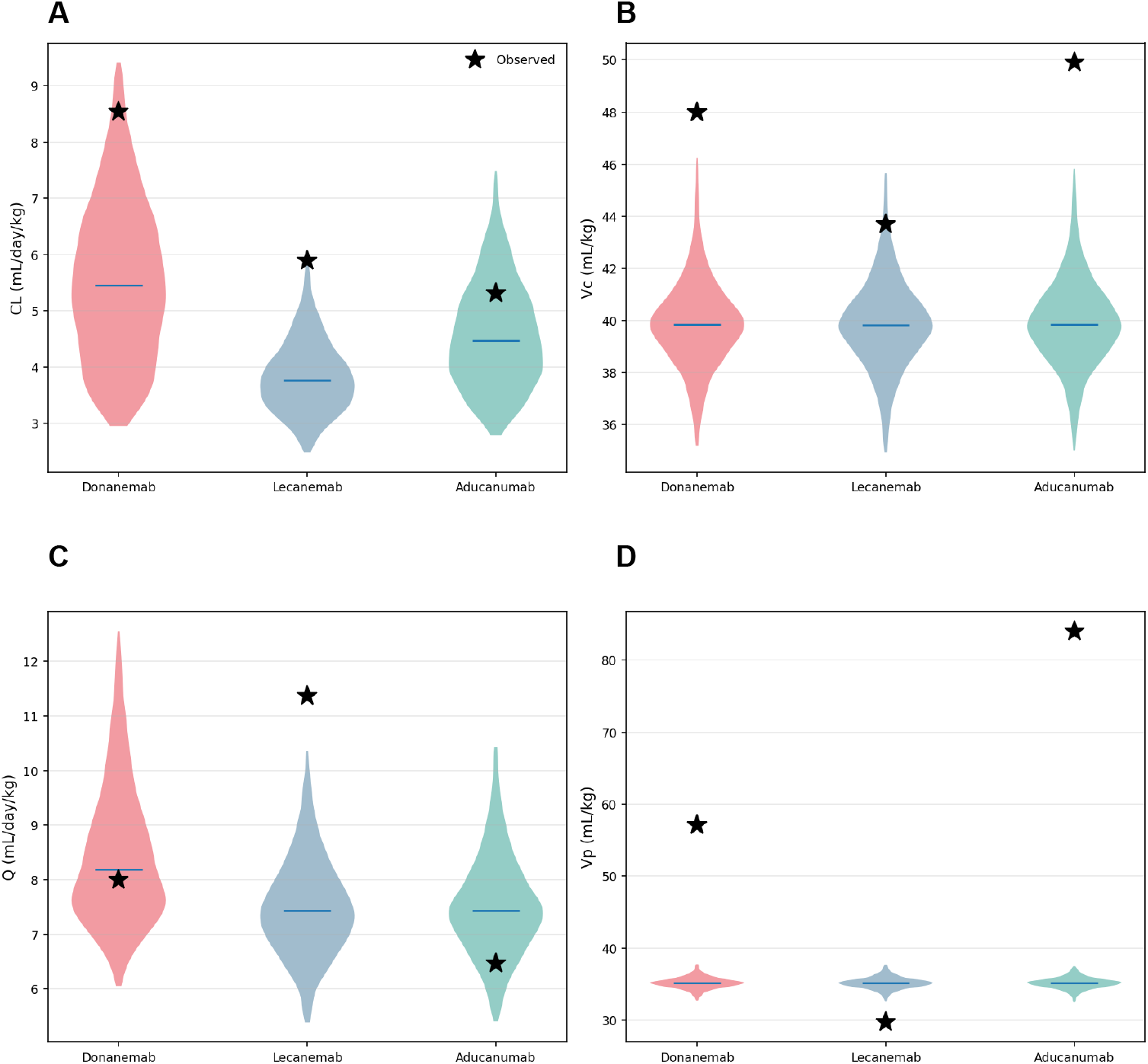
Posterior predictive distributions of human two-compartment PK parameters for each CNS-targeted anti-amyloid mAb. Violin plots showing the posterior predictive distributions of clearance (CL, mL/day/kg; A), central volume of distribution (Vc, mL/kg; B), inter-compartmental clearance (Q, mL/day/kg; C), and peripheral volume of distribution (Vp, mL/kg; D) for donanemab (red), lecanemab (blue), and aducanumab (teal). Horizontal blue lines within each violin indicate the posterior predictive median. Black stars denote the observed human values from published population PK analyses. For CL, observed values (stars) fall above the posterior predictive distributions for all three mAbs, consistent with target-mediated clearance not captured by the model. For Vc, predicted medians (∼39.8 mL/kg) were reasonably consistent with observed values (48.0 mL/kg for donanemab, 43.7 mL/kg for lecanemab, 49.9 mL/kg for aducanumab). For Q, predicted and observed values show reasonable agreement. For Vp, aducanumab shows a notably large observed value (84.0 mL/kg, from V2 = 6.04 L / 71.9 kg) substantially exceeding the predicted distribution (∼35 mL/kg), suggesting unusually extensive tissue distribution for this antibody.

For clearance (CL), observed values (black stars) fell above the posterior predictive distributions for all three mAbs, consistent with the TMDD-driven under-prediction. For central volume of distribution (Vc), predicted medians (∼39.8 mL/kg) were reasonably consistent with observed values (48.0 mL/kg for donanemab, 43.7 mL/kg for lecanemab, 49.9 mL/kg for aducanumab), reflecting the population-level prior learned from the training set. For peripheral volume (Vp), aducanumab was a notable outlier: observed Vp = 84.0 mL/kg (from V2 = 6.04 L / 71.9 kg) was substantially larger than predicted (∼35 mL/kg), suggesting unusually extensive tissue distribution for this antibody.

## 3. DISCUSSION

In this study, we developed and validated a Bayesian hierarchical PK framework for predicting human pharmacokinetics and recommending FIH doses for therapeutic mAbs, with emphasis on uncertainty quantification. While Bayesian methods have been widely applied in population PK modeling and dose individualization, their use for propagating uncertainty through allometric scaling in mAb FIH dose selection has not been systematically explored. The framework was calibrated on two-compartment PK data from 9 training mAbs and achieved a MAPE of 11.6% in leave-one-out cross-validation, with all 9 predictions within 2-fold of observed values. To evaluate the framework in a clinically demanding setting, the calibrated model was applied to three CNS-targeting mAbs approved for Alzheimer’s disease, donanemab, lecanemab, and aducanumab, using only their cynomolgus PK data. All three prospective-style predictions fell within the 2-fold boundary, demonstrating that the Bayesian framework provides accurate and uncertainty-aware FIH dose recommendations.

A key advantage of the Bayesian approach over conventional allometric scaling is the provision of full posterior predictive distributions rather than point estimates. For FIH dose selection, this enables explicit risk quantification to evaluate the probability that a candidate dose will exceed a safety threshold (e.g., the probability that C_max_ exceeds a level associated with ARIA) or fall below an efficacy target (e.g., the probability that brain exposure is insufficient for target engagement). The 90% credible intervals reported here (e.g., CL of 3.16 – 6.34 mL/day/kg for aducanumab, 3.38 – 8.17 mL/day/kg for donanemab) provide a natural framework for such assessments and are directly interpretable.

The MABEL-based dose estimation incorporated a brain-to-plasma ratio of 0.2%, consistent with the literature on IgG blood-brain barrier (BBB) penetration [11]. This is a critical consideration for CNS-targeted mAbs, where the pharmacologically relevant exposure is in the brain and the orders-of-magnitude attenuation across the BBB must be explicitly modeled. The framework’s ability to propagate PK parameter uncertainty through the brain exposure calculation ensures that the MABEL estimate reflects cumulative uncertainty in CL, Vc, and the BBB penetration ratio.

Several features of the Bayesian hierarchical framework distinguish it from conventional allometric approaches. First, the precision-weighted combination of population-level priors and allometric predictions naturally adapts to the informativeness of each source when the allometric scaling is precise (small *σ*_*scale*_), the prediction is dominated by the scaled cynomolgus CL; when inter-mAb variability is small (small *σ*_*CL*_), the population prior exerts greater influence. This adaptive weighting is particularly valuable for mAbs with atypical cynomolgus PK, where pure allometric scaling might yield unreliable predictions. Second, the hierarchical structure provides partial pooling across the training mAbs, which protects against overfitting to individual antibodies and yields more robust population-level estimates than fitting each mAb independently. Third, the framework is extensible as new mAbs with known human PK become available, the training set can be expanded and the model re-fit, progressively refining the population priors and scaling relationship. Fourth, the framework is broadly applicable to any therapeutic mAb. The population priors and allometric relationship are not specific to a particular target class or indication, and the CNS test case presented here demonstrates its utility in a high-stakes setting where uncertainty quantification is most valuable.

Several limitations should be considered when interpreting these results. First, the training set comprised 9 mAbs, which limits the precision of the population-level hyperparameters and may not fully represent the diversity of mAb PK behavior. The training set was composed predominantly of IgG1 antibodies targeting peripheral antigens, and the generalizability to other IgG subclasses (e.g., IgG4) or antibodies with unusual PK properties (e.g., recycling antibodies, bispecifics) requires further evaluation. Second, the two-compartment linear PK model does not account for TMDD, which proved to be the dominant source of prediction error for the CNS test mAbs. Incorporating a quasi-steady-state or Michaelis-Menten TMDD component into the hierarchical framework could improve predictions for mAbs with significant target-mediated clearance, though this would require target expression and binding affinity data that may not be available at the FIH stage. Third, volume and inter-compartmental clearance parameters for some training mAbs were estimated from typical mAb values rather than directly measured from two-compartment population PK analyses, which introduces additional uncertainty not fully captured in the model. Fourth, the MABEL calculation relied on a fixed brain-to-plasma ratio of 0.2% for all three test mAbs, whereas in practice this ratio may vary with dose, disease state, and BBB integrity in Alzheimer’s disease patients [37]. Fifth, the retrospective validation was limited to three mAbs, all targeting amyloid-beta. Prospective application to a broader panel of CNS-targeted biologics including those targeting tau, alpha-synuclein, or neuroinflammatory pathways as well as mAbs for other indications with narrow therapeutic windows, would strengthen the evidence for the framework’s generalizability. Finally, the model assumes that the allometric scaling relationship learned from peripheral mAbs is directly transferable to CNS-targeted mAbs. While systemic PK is likely similar (all are IgG1 molecules cleared primarily through FcRn-mediated recycling and catabolism), the relevance of CNS target engagement to systemic clearance particularly for mAbs with high target burden in the brain remains an area of active investigation.

In conclusion, the Bayesian hierarchical PK framework presented here provides a principled, uncertainty-aware approach to FIH dose selection for therapeutic mAbs. The framework achieved high predictive accuracy in cross-validation (MAPE 11.6%) and generated clinically appropriate FIH dose recommendations for all three test anti-amyloid antibodies, with prediction errors of -36.1%, -36.1%, and -15.7% for donanemab, lecanemab, and aducanumab, respectively. The systematic under-prediction of clearance for mAbs with significant TMDD is shared by all allometric scaling methods and represents a pharmacologically conservative bias (over-prediction of exposure, wider safety margins). This limitation could be addressed in future work by incorporating target-mediated clearance components. The framework is broadly applicable to therapeutic mAbs and is particularly well-suited to settings where the therapeutic window is narrow such as CNS-targeted biologics, where the margin between efficacy (requiring sufficient brain exposure across the BBB) and safety (ARIA risk) demands rigorous uncertainty quantification at the FIH dose selection stage.

## 5. AUTHOR CONTRIBUTIONS

BR and AG conceived and designed the study. Both authors contributed to revisions across all sections and approved the final manuscript.

## REFERENCES

1. Wang W, Wang E, Balthasar J. Monoclonal antibody pharmacokinetics and pharmacodynamics. Clinical Pharmacology & Therapeutics 2008;84:548–58.

2. Deng R, Iyer S, Theil F-P, Mortensen DL, Fielder PJ, Prabhu S. Projecting human pharmacokinetics of therapeutic antibodies from nonclinical data: what have we learned? vol. 3, Taylor & Francis; 2011, p. 61–6.

3. Gelman A, Carlin JB, Stern HS, Rubin DB. Bayesian data analysis. Chapman and Hall/CRC; 1995.

4. Wakefield J. Bayesian and frequentist regression methods. vol. 23. Springer; 2013.

5. Schmidli H, Gsteiger S, Roychoudhury S, O’Hagan A, Spiegelhalter D, Neuenschwander B. Robust meta-analytic-predictive priors in clinical trials with historical control information. Biometrics 2014;70:1023–32.

6. Neuenschwander B, Branson M, Gsponer T. Critical aspects of the Bayesian approach to phase I cancer trials. Statistics in Medicine 2008;27:2420–39.

7. Wang X, Suttner L, Jemielita T, Li X. Propensity score-integrated Bayesian prior approaches for augmented control designs: a simulation study. Journal of Biopharmaceutical Statistics 2022;32:170–90.

8. Han L, Deng Q, He Z, Fleischer F, Yu F. Bayesian hierarchical model for dose-finding trial incorporating historical data. Journal of Biopharmaceutical Statistics 2024;34:646–60.

9. Haraya K, Kuramochi T. Quantitative Prediction of Human Pharmacokinetics for Fc-Engineered Therapeutic Monoclonal Antibodies With Increased FcRn Binding Mutations After Subcutaneous Injection. Clinical and Translational Science 2026;19:e70482.

10. Chang H-Y, Wu S, Meno-Tetang G, Shah DK. A translational platform PBPK model for antibody disposition in the brain. Journal of Pharmacokinetics and Pharmacodynamics 2019;46:319–38.

11. Poduslo JF, Curran GL, Berg CT. Macromolecular permeability across the blood-nerve and blood-brain barriers. Proceedings of the National Academy of Sciences 1994;91:5705–9.

12. Sperling RA, Jack Jr CR, Black SE, Frosch MP, Greenberg SM, Hyman BT, et al. Amyloid-related imaging abnormalities in amyloid-modifying therapeutic trials: recommendations from the Alzheimer’s Association Research Roundtable Workgroup. Alzheimer’s & Dementia 2011;7:367–85.

13. Pardridge WM. Blood-brain barrier and delivery of protein and gene therapeutics to brain. Frontiers in Aging Neuroscience 2020;11:373.

14. Skarlis C, Angelopoulou E, Rentzos M, Papageorgiou SG, Anagnostouli M. Monoclonal Antibodies as Therapeutic Agents in Autoimmune and Neurodegenerative Diseases of the Central Nervous System: Current Evidence on Molecular Mechanisms and Future Directions. International Journal of Molecular Sciences 2025;26:9398.

15. Sims JR, Zimmer JA, Evans CD, Lu M, Ardayfio P, Sparks J, et al. Donanemab in early symptomatic Alzheimer disease: the TRAILBLAZER-ALZ 2 randomized clinical trial. Jama 2023;330:512–27.

16. Van Dyck CH, Swanson CJ, Aisen P, Bateman RJ, Chen C, Gee M, et al. Lecanemab in early Alzheimer’s disease. New England Journal of Medicine 2023;388:9–21.

17. Budd Haeberlein S, Aisen PS, Barkhof F, Chalkias S, Chen T, Cohen S, et al. Two randomized phase 3 studies of aducanumab in early Alzheimer’s disease. The Journal of Prevention of Alzheimer’s Disease 2022;9:197–210.

18. Lu J-F, Bruno R, Eppler S, Novotny W, Lum B, Gaudreault J. Clinical pharmacokinetics of bevacizumab in patients with solid tumors. Cancer Chemotherapy and Pharmacology 2008;62:779–86.

19. Garg A, Quartino A, Li J, Jin J, Wada DR, Li H, et al. Population pharmacokinetic and covariate analysis of pertuzumab, a HER2-targeted monoclonal antibody, and evaluation of a fixed, non-weight-based dose in patients with a variety of solid tumors. Cancer Chemotherapy and Pharmacology 2014;74:819–29.

20. Cosson VF, Ng VW, Lehle M, Lum BL. Population pharmacokinetics and exposure–response analyses of trastuzumab in patients with advanced gastric or gastroesophageal junction cancer. Cancer Chemotherapy and Pharmacology 2014;73:737–47.

21. Hayashi N, Tsukamoto Y, Sallas WM, Lowe PJ. A mechanism-based binding model for the population pharmacokinetics and pharmacodynamics of omalizumab. British Journal of Clinical Pharmacology 2007;63:548–61.

22. U.S. Food and Drug Administration. BLA 103792. Trastuzumab (Herceptin), 2012.

23. U.S. Food and Drug Administration. BLA 125085. Bevacizumab (Avastin), 2006.

24. U.S. Food and Drug Administration. BLA 125409. Pertuzumab (Perjeta), 2011.

25. U.S. Food and Drug Administration. BLA 103976. Omalizumab (Xolair), 2003.

26. U.S. Food and Drug Administration. BLA 761248. Donanemab (Kisunla), 2023.

27. U.S. Food and Drug Administration. BLA 761269. Lecanemab (Leqembi), 2023.

28. U.S. Food and Drug Administration. BLA 761178. Aducanumab (Aduhelm), 2023.

29. Gueorguieva I, Willis BA, Chua L, Chow K, Ernest CS, Shcherbinin S, et al. Donanemab population pharmacokinetics, amyloid plaque reduction, and safety in participants with Alzheimer’s disease. Clinical Pharmacology & Therapeutics 2023;113:1258–67.

30. Hayato S, Takenaka O, Sreerama Reddy SH, Landry I, Reyderman L, Koyama A, et al. Population pharmacokinetic-pharmacodynamic analyses of amyloid positron emission tomography and plasma biomarkers for lecanemab in subjects with early Alzheimer’s disease. CPT: Pharmacometrics & Systems Pharmacology 2022;11:1578–91.

31. Kandadi Muralidharan K, Tong X, Kowalski KG, Rajagovindan R, Lin L, Budd Haberlain S, et al. Population pharmacokinetics and standard uptake value ratio of aducanumab, an amyloid plaque–removing agent, in patients with Alzheimer’s disease. CPT: Pharmacometrics & Systems Pharmacology 2022;11:7–19.

32. Dormand JR, Prince PJ. A family of embedded Runge-Kutta formulae. Journal of Computational and Applied Mathematics 1980;6:19–26.

33. Hoffman MD, Gelman A. The No-U-Turn sampler: adaptively setting path lengths in Hamiltonian Monte Carlo. J Mach Learn Res 2014;15:1593–623.

34. Gelman A, Hill J. Data analysis using regression and multilevel/hierarchical models. Cambridge university press; 2007.

35. Logovinsky V, Satlin A, Lai R, Swanson C, Kaplow J, Osswald G, et al. Safety and tolerability of BAN2401-a clinical study in Alzheimer’s disease with a protofibril selective Aβ antibody. Alzheimer’s Research & Therapy 2016;8:14.

36. Sims JR, Zimmer JA, Evans CD, Lu M, Ardayfio P, Sparks J, et al. Donanemab in early symptomatic Alzheimer disease: the TRAILBLAZER-ALZ 2 randomized clinical trial. Jama 2023;330:512–27.

37. Zlokovic BV. The blood-brain barrier in health and chronic neurodegenerative disorders. Neuron 2008;57:178–201.

